# Cloud-enabled Scalable Analysis of Large Proteomics Cohorts

**DOI:** 10.1101/2024.09.05.611509

**Authors:** Harendra Guturu, Andrew Nichols, Lee S. Cantrell, Seth Just, Janos Kis, Theodore Platt, Iman Mohtashemi, Jian Wang, Serafim Batzoglou

## Abstract

Rapid advances in depth and throughput of untargeted mass-spectrometry-based proteomic technologies are enabling large-scale cohort proteomic and proteogenomic analyses. As such studies scale, the data infrastructure and search engines required to process data must also scale. This challenge is amplified in search engines that rely on library-free match between runs (MBR) search, which enable enhanced depth-per-sample and data completeness. However, to-date, no MBR-based search could scale to process cohorts of thousands or more individuals. Here, we present a strategy to deploy search engines in a distributed cloud environment without source code modification, thereby enhancing resource scalability and throughput. Additionally, we present an algorithm, Scalable MBR, that replicates the MBR procedure of the popular DIA-NN software for scalability to thousands of samples. We demonstrate that Scalable MBR can search thousands of MS raw files in a few hours compared to days required for the original DIA-NN MBR procedure and demonstrate that the results are almost indistinguishable to those of DIA-NN native MBR. The method has been tested to scale to over 15,000 injections and is available for use in the Proteograph™ Analysis Suite.

## Introduction

Mass spectrometry (MS) based proteomics has revolutionized the field of protein analysis, standing as a cornerstone technology for detecting thousands of proteins within a single sample. MS proteomics enables the rapid, robust, and sensitive detection of a broad range of proteins, allowing for the comprehensive profiling of complex biological samples.^1,2^ Recent advances in instrumentation— particularly in speed, specificity, and detector sensitivity—have enabled plasma proteomics to achieve unprecedented depth and throughput.^3,4^ With these evolving technologies, biological insights have accelerated, including in cultured- and single-cell measurements as reviewed by Peters-Clark and colleagues.^5^ Simultaneously, enrichment methods for blood plasma measurements have significantly improved proteome depth compared to untreated plasma and depletion methods.^6,7^ These advancements allow for the scaling cohort studies from tens of samples^8,9^ to thousands.^6,10^ As proteomic cohort measurements continue to expand, studies involving tens to hundreds of thousands of samples, comparable to the scale of current genomic studies, are becoming increasingly feasible.^11^ The added scale and depth provides researchers with the statistical power necessary to generate meaningful insights for diagnostics or other clinically significant outcomes, as evidenced by Bueno Álvez and colleagues’ study involving over 1,400 samples, which identified multiple cancer biomarkers^12^, or Koh and colleagues’ identification of lung cancer biomarkers from over 2,500 samples.^13^

Mass spectrometers have recently undergone significant improvements in throughput.^3,14^ Utilizing untargeted data-independent acquisition (DIA) methods, throughput of up to 300 samples per day has been reported for whole proteome measurements.^3^ However, a concurrent progression in scalable proteomics search engines has lagged behind. Current software solutions are challenged by the massive (terabyte scale) datasets generated by faster, more sensitive instruments. It is often difficult, if not impossible, to effectively search more than 500 sample file outputs from MS studies on a single-node workstation.^15,16^

In the context of growing cohort sizes and single-node limitations, cloud-based distributed computing infrastructure has emerged as an attractive solution. Leveraging the cloud allows for the distribution of computational tasks across multiple servers, facilitating more extensive and in-depth cohort-level studies.^17^ Cloud computing for software scalability in proteomics data processing represents a crucial shift for maintaining momentum in discovery proteomics. Workflow orchestration engines such as NextFlow and Galaxy framework have indeed been proposed for implementing such data processing in the past.^18,19^ However, the cloud engineering expertise required to configure the supporting cloud infrastructure typically limits adoption of these frameworks.

In addition to a highly parallelized implementation of the popular DIA search engine DIA-NN^20^(Data-Independent Analysis using Neural Networks), we present Scalable MBR, a new method for extending DIA-NN’s match between runs (MBR) strategy to large cohorts. Scalable MBR maximizes a study’s sensitivity and power by increasing data completeness compared to standard library free search (**SFigure 1**). We leverage cloud-computing resources, including AWS, and orchestrate pipelines using Prefect Cloud to deploy the pipeline in the Proteograph™ Analysis Suite (PAS) for a user-friendly experience.^21^

## Methods

### Orchestration of distributed pipelines in the cloud

The MapReduce programming model is used to scale up the search of MS raw files.^22^ In brief, the programming model “**maps**” each MS raw file (large - approximately several gigabytes) to an independent node to search for peptide-spectrum matches. Since each MS raw file is searched independently, this step can be parallelized up to the number of available nodes and additional files can be queued to run as the jobs finish. In the “**reduce**” step, the intermediate search results (small - on the order of several megabytes) are gathered onto a single node to perform protein inference and global false discovery rate (FDR) calculations. Adding MBR to this paradigm is challenging due to the need to share information across all files. As implemented in DIA-NN, this requires the single node at the reduce step to access all raw MS files. This requirement limits scalability to locally available disk space and adds significant overhead for file transfer. This bottleneck can be bypassed with techniques such as “ID-RT-IM profiling”, which produce semi-empirical libraries with only intermediate result files^23^ but affect the accuracy and consistency of identification and quantification. Ensuring the availability of sufficient storage in a cloud execution environment also poses difficulties in configuring appropriate resource limits to optimize the tradeoff between peak scalability and cost efficiency. This is further complicated when using containerized execution environments, in which multiple jobs can share a single underlying compute node to minimize costs, as resource limits must be chosen to account for all jobs that might run concurrently on a single node. More flexible network attached storage solutions exist, but those often compromise IO performance which is critical for processing large raw MS files.

### Extending the distributed pipeline with a Scalable MBR strategy

The Scalable MBR pipeline, summarized in **Figure 1**, extends the parallel search pipeline. When parallel searches (green blocks) are run for each input file, to avoid bottlenecking library generation of MBR on the “reduce” step, empirical libraries are generated for each MS raw file during the many “map” steps. Then, during the “reduce” step (purple), DIA-NN is allowed to process all the individual quant files as usual to generate a report with global FDR. For each identified precursor, the report is employed to select the MS raw file with the lowest global precursor FDR and the precursor spectrum is extracted from the raw file’s corresponding empirical library. All extracted empirical precursor spectra are merged into a final “**Scalable MBR**” library. Importantly, this process requires only small intermediate empirical library and search result files. While ultimate scalability is still limited by storage requirements, they are massively reduced, allowing for the processing of very large datasets with limited resources and simplifying the configuration of cost-effective cloud infrastructure. The final library is used in another distributed map-reduce process (orange and blue blocks in figure 1) to generate final MBR results. This approach approximates DIA-NN’s empirical library building and native MBR with `--smart-profiling` and is an alternative to using a semi-empirical library (ID-RT-IM profiling) in the second pass.

**Figure 1:**
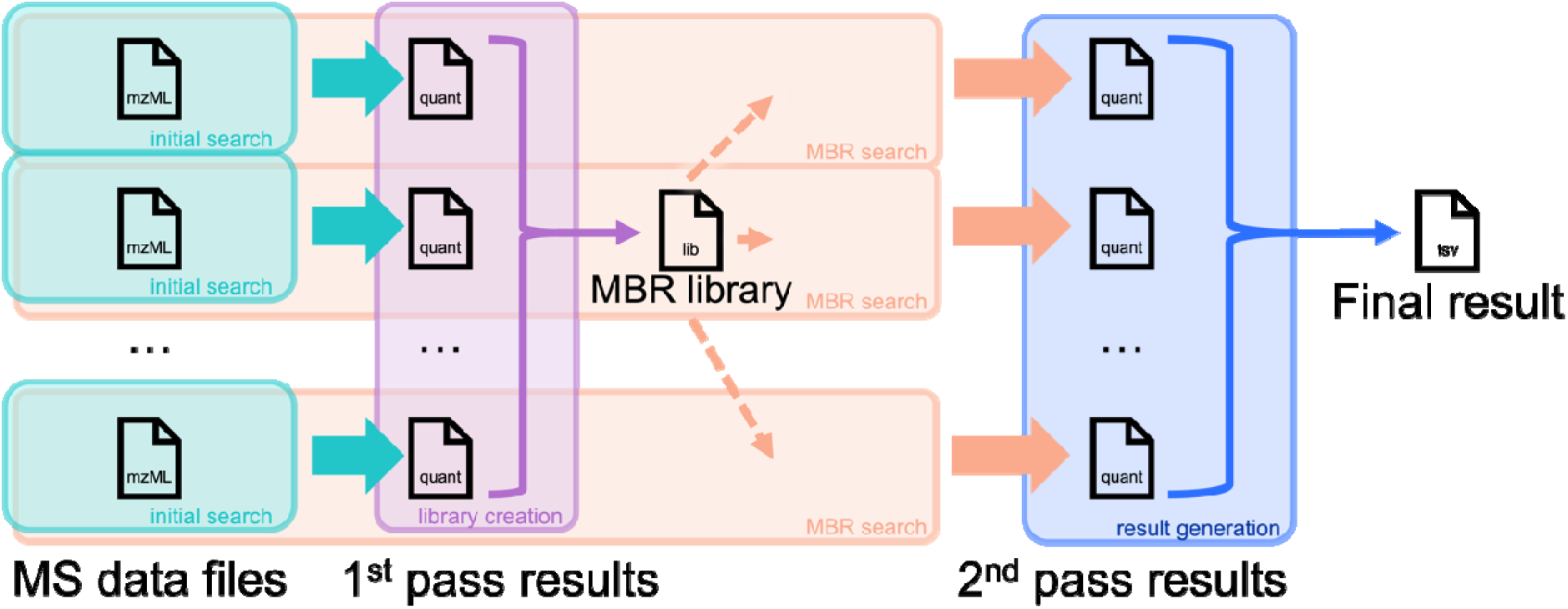
Overview of scalable cloud-enabled architecture with match between runs. Parallel searches of individual input files (green blocks) generate intermediate results files that are processed on single node (purple) to generate an empirical MBR library. Subsequent parallel searches (orange) generate per-file results which are processed (blue) to give a final MBR result for the whole experiment. Importantly, only parallelized map steps (green/orange) require access to raw data, while global reduce steps (purple/blue) use only small results files, permitting scalability to thousands of files with modest compute and storage requirements for all processing nodes.

### Dataset for evaluation of Scalable MBR

The evaluation of scalability and concordance of Scalable MBR to DIA-NN’s MBR was done across a range of cohort acquires on timsTOF, Orbitrap™ Exploris™ and Astral™ analyzers.^10,24–28^ Since many of these cohorts are currently being analyzed for biological insights and have not been publicly released, we limited the analyses in this work to one that has been deposited at PXD042852.^24^ The cohort is composed of 1,840 MS raw files acquired on Bruker timsTOF Pro2 using dia-PASEF with a 22 minute gradient and a 33 minute total runtime per sample.^14^

The files were all searched in library free mode with the following common DIA-NN parameters:

~~~
--mass-acc-ms1 10, --mass-acc 10, --qvalue 0.1, --matrices, --met-
excision, --cut K*,R*, --relaxed-prot-inf, --reannotate, --threads 32,
--predictor, --unimod4, --use-quant, --peak-center, --no-ifs-removal
~~~

The following MBR strategies were used to compare with Scalable MBR.

- DIA-NN MBR: `--smart-profiling --reanalyze`
  - Empirical library building and second pass on one node/process.
  - The baseline strategy to match.
- DIA-NN MBR Distributed Pass`: --smart-profiling`
  - Empirical library building on one node, but second pass is manually distributed.
  - An alternate baseline that shows the differences contributed by doing the second pass in separate process (due to differences such as slightly different decoys for FDR scoring).
- ID-RT-IM MBR: `--rt-profiling --reanalyze`
  - Semi-empirical library used in second pass on one node/process.
  - This is an alternative to avoid the large bottleneck of requiring all MS raw files on one node for empirical library building, but limits user to theoretical spectra with empirical ion-mobility and retention time values.
- ID-RT-IM MBR Distributed Pass: `--rt-profiling`
  - Semi-empirical library built on one node, and second pass is manually distributed.
  - Included for completeness with the other comparisons.
- Scalable MBR:
  - The method introduced in this work delivers an empirical library while avoiding the bottleneck of requiring all MS raw files on one node and automatically distributes the 2^nd^ pass.

For the evaluation, the metrics of success were identification rate (how well do alternate strategies recover the precursors and protein groups), quantification (how similar is the quantification) and total run time compared to the baseline DIA-NN MBR.

### System Provisioning for Runtime and Cost Estimation

Single node DIA-NN MBR was performed on an AMD 16-core, 128 GB AWS EC2 instance with SSD attached. Instance configuration was optimized for cost-to-performance and represents a best-case scenario for up to 1,000 files. Cost was based on spot pricing for EC2 instances and price-per-hour of storage. Additional costs associated with S3 storage, transfer, and long-term storage provisioning were not included.

Cloud pipeline DIA-NN with or without MBR is run in two different steps using AWS Batch, with provisioning tailored to suit the needs of the respective step.

The map step to search each raw MS file is an embarrassingly parallel process and is run with provisioning based on the library file size. This has been found to be the single greatest factor impacting the amount of memory required to execute the step. Each data file is run in a separate Docker container spot instance that has been configured to execute the DIA-NN algorithm using the Ubuntu 22.04 operating system. For libraries under 1 GB in size, provisioning the instance with 8 vCPU and 32 GB RAM will typically suffice. Larger libraries are run with 16 vCPU and 64 GB RAM. Using the least amount of resources is imperative at this step if cost is a concern. Since this step is performed in parallel, any cost due to over-provisioning resources will be multiplied n-fold vis-à-vis the number of containers used,

In the second step, to perform FDR and protein inference, all search result files and empirical library files when using Scalable MBR files are downloaded into a single container and DIA-NN is run with the “--use-quant” flag. The container used in this step is typically provisioned with 32 vCPU and 480 GB RAM and a 5TB SSD hard drive. Since this process is only run on a single instance, overprovisioning to this extent has a negligible effect on cost.

## Results and Discussion

The performance of different search regimes was evaluated on a dataset of 1,840 MS raw files. To demonstrate the utility of MBR, a comparison of DIA-NN library free search and DIA-NN MBR was performed. Increased depth, data completeness, and data similarity across the cohort were observed to improve with MBR, enhancing possible downstream cohort analyses (**SFigure 1**). Library-free search enables minimally biased search; however, it is unable to detect precursor features of lower intensity in some samples within a biologically heterogenous cohort. MBR addresses this limitation by first performing a library free search and then filtering the predicted library on empirically observed precursor features, augmenting experimental spectral intensities, retention time and ion mobility with empirical observations. Prior studies have also supported the benefit of reducing library size for search sensitivity^29^ and supported the use of empirical spectral attributes for enhanced data quality.^30^

The performance of Scalable MBR relative to DIANN MBR and three other approaches (see methods) was evaluated for consistency in qualitative identification and quantitative inference. Comparison between Scalable MBR and DIA-NN MBR showed similar depth and significant overlap, >90%, of precursor and protein groups between algorithms (**Figure 2A)**. Each search method showed an approximately equal count of unique features. Similar comparison of distinct and common features is available when comparing DIA-NN MBR to alternate strategies (see methods) in **SFigure 2**. Notably, the distribution of unique and common precursors in DIA-NN MBR Distributed Pass and DIA-NN MBR is comparable to that of Scalable MBR and DIA-NN MBR. This suggests that the difference is primarily being driven due the distribution process rather than the quality of the library generated by Scalable MBR since DIA-NN MBR Distributed Pass uses the baseline library generated by DIA-NN MBR. Chipset instructions and exact library composition may have similar impact on minor qualitative ID inconsistency between approaches. Additionally, Scalable MBR demonstrated more consistent identifications to DIA-NN MBR compared to ID-RT-IM-based MBR, supporting the value of empirically derived spectra in measurements. The comparison with the “Distributed Pass” searches that manually use the derived library for distributed second pass result in different protein groups from DIA-NN MBR which is not desired. Scalable MBR correctly handles the protein grouping in the second pass to match DIA-NN MBR. Moreover, the distributed pass by Scalable MBR provides an additional speedup, since typically DIA-NN and other implementations of MBR would continue the second pass search of the raw files using the library in the same process which again requires long wait times when there are thousands of files since each file is processed sequentially.^23^

**Figure 2:**
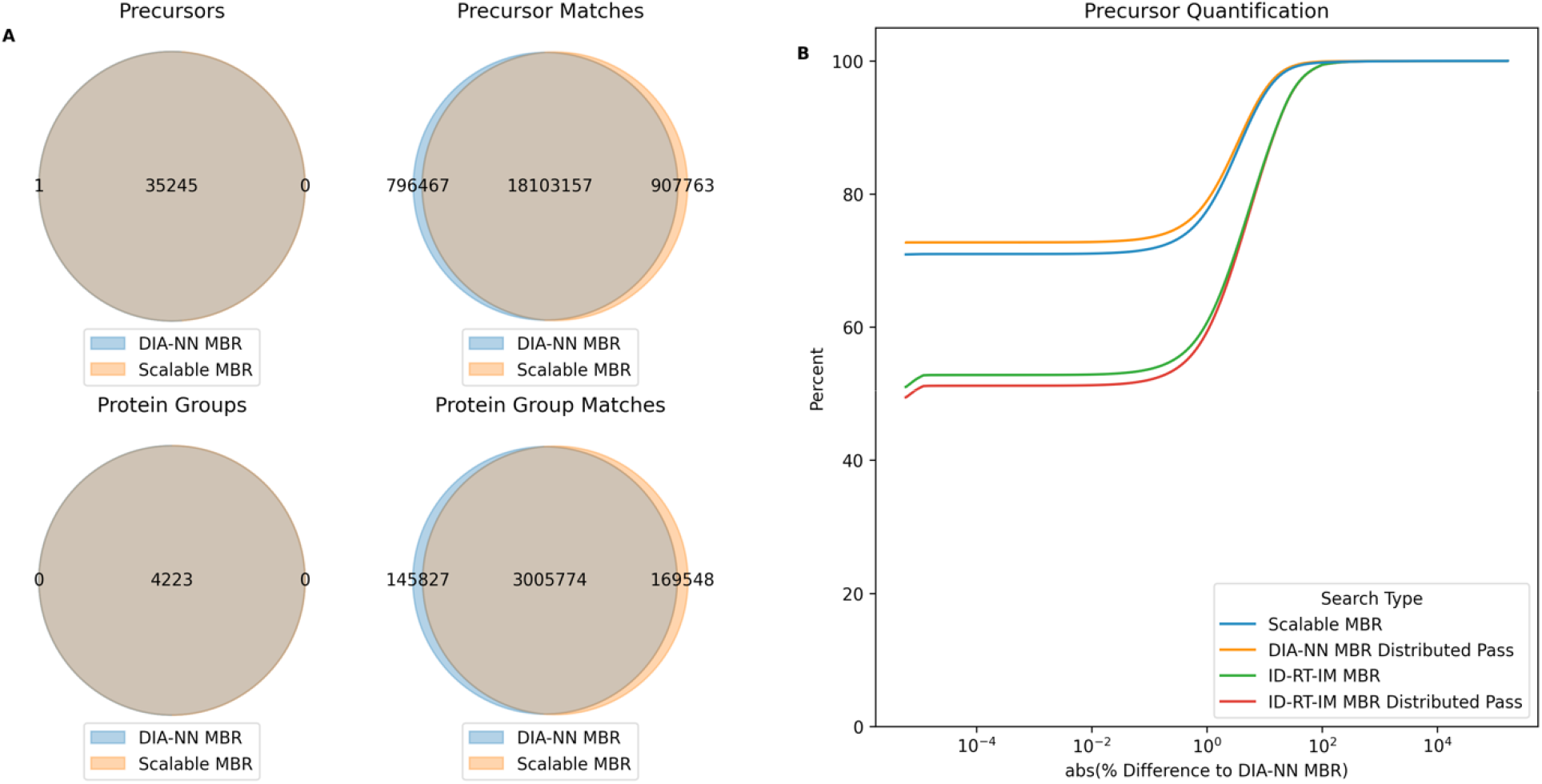
Comparison of cloud-enabled MBR strategies to DIA-NN’s default single node MBR. **A)** Scalable MBR identifies essentially the same number of unique precursors and protein groups. Further, the total number of matches are largely common and of similar count. **B)** The similarity of precursor quantification is compared to DIA-NN’s MBR for Scalable MBR and a few alternative options. The library from Scalable MBR and DIA-NN MBR Distributed Pass (where the library building is done on a single node by DIA-NN, but the second pass is distributed) have essentially identical quantification performance.

To evaluate consistency in quantification, the percent absolute difference of the quantification value with the one from DIA-NN MBR was computed and the accumulation curves of difference are shown in **Figure 2B** for precursors and in **SFigure 3** for protein groups. As can be seen, Scalable MBR and DIA-NN MBR Distributed Pass show very similar quantification differences and over 70% of the matches have near identical quantification and essentially all matches are within 10% difference. DIA-NN MBR Distributed Pass is the theorical best approximation of DIA-NN MBR with distribution and Scalable MBR which is also distributed performs very similarly. Additionally, we see that ID-RT-IM based strategies show lower concordance in quantification with DIA-NN MBR.

Finally, we benchmark the runtime and cost of Scalable MBR compared to DIA-NN MBR. **Figure 3A**, shows that for small datasets (<30 files) for which a desktop application might be preferable, the run time is comparable, but separation grows rapidly, and the runtime is untenable at the scale of 1000 files where DIA-NN MBR would take nearly a week compared to a few hours with Scalable MBR. Furthermore, we show in **Figure 3B**, that although Scalable MBR delivers large speed ups, the costs are comparable to DIA-NN MBR (since in the cloud parallel compute and serial compute cost the same).

**Figure 3:**
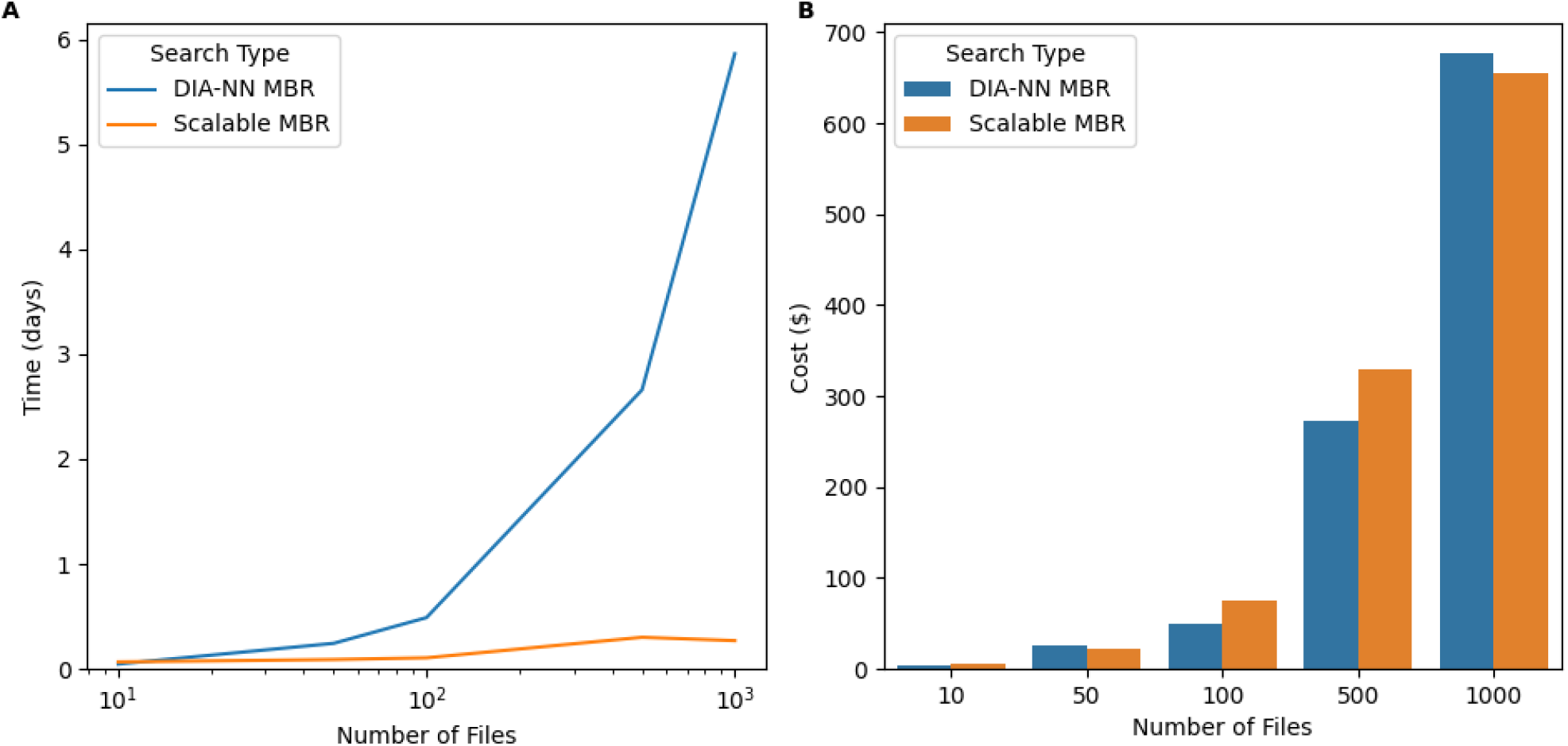
Scalable MBR maintains similar cost to DIA-NN while returning results orders of magnitude faster for large cohorts. **A)** Scalable MBR starts returning results noticeable faster than the single node MBR strategy even for cohorts with as few as 100 files. The difference compounds for larger cohorts. **B)** The cost between Scalable MBR and DIA-NN MBR is essentially the same (since distribution only parallelizes the cost accumulation).

To further evaluate practical limits of Scalable MBR and measurable depth of the plasma proteome, a set of 7,200 human plasma and serum sample enriched using Proteograph XT and analyzed on Orbitrap Astral were searched, in total exceeding 1.6 billion spectra. For these files, Scalable MBR library compilation, distributed search and report generation completed in 30 hours to identify more than 10,000 plasma protein groups, shared in the Seer Protein Discovery Catalog.^31^ A significant portion of search time was spent on MaxLFQ protein intensity calculation which scales quadratically with sample count.^32,33^ Nonetheless, Scalable MBR was successfully extended to complete a search at an extrapolated 30x faster than DIA-NN MBR while simultaneously highlighting a key value of MS data acquisition: that datasets may be re-processed with new libraries or other search modifications to improve statistical inference.

## Conclusion

In this work, we present a framework to search large (hundreds to thousands) DIA MS proteomic datasets seamlessly in the cloud. We further extend the framework with an approximate algorithm for smart profiling of empirical spectral libraries called Scalable MBR. This Scalable MBR algorithm matches the performance of DIA-NN’s built in MBR with smart profiling without the bottleneck of requiring a large server with terabytes of disk space to consolidate all the MS raw files and further spending days to weeks searching the datasets.

This allows MS proteomics users to rapidly search and re-search their data to identify biological insights while benefiting from the increased data completeness of MBR to maximize sensitivity and power in large cohort analyses. This framework has been used in multiple large cohort studies that are currently in various stages of analysis, peer review and publication.^10,24–28,34^ We have incorporated this algorithm into the Proteograph™ Analysis Suite (PAS) for users to search their large cohorts without any cloud deployment expertise. This work lowers the barrier for large cohort MS proteomics and enables new biological discoveries due to the unlocked scale.

## Author Statements

### Data Availability

The raw MS-proteomics data used in this study is available via ProteomeXchange with identifier PXD042852 (https://proteomecentral.proteomexchange.org/cgi/GetDataset?ID=PXD042852).

## Acknowledgements

We are grateful to Karsten Suhre *et al*. and Weill Cornell Medicine - Qatar for releasing their data at PXD042852 as part of their study to enable this work. We also want to thank Amir Alavi, Ting Huang, Benjamin Lacar, Alexey Stukalov, Guhan R. Venkataraman, & Jian Yao for incorporating the Scalable MBR method into their studies to help evaluate the method’s performance.

## Author Contributions

Study Design: H.G., J.W., S.B.

Implementation: H.G., A.N., J.K., T.P., I.M.

Data Analysis: H.G., L.S.C., S.J., J.W.

Manuscript Writing: H.G., L.S.C, S.J., S.B.

All authors contributed to critically reviewing the results and the manuscript.

## Competing Interests

All authors are employees and/or stockholders of Seer, Inc.

