## Supplementary Information for "Cloud-enabled Scalable Analysis of Large Proteomics Cohorts"

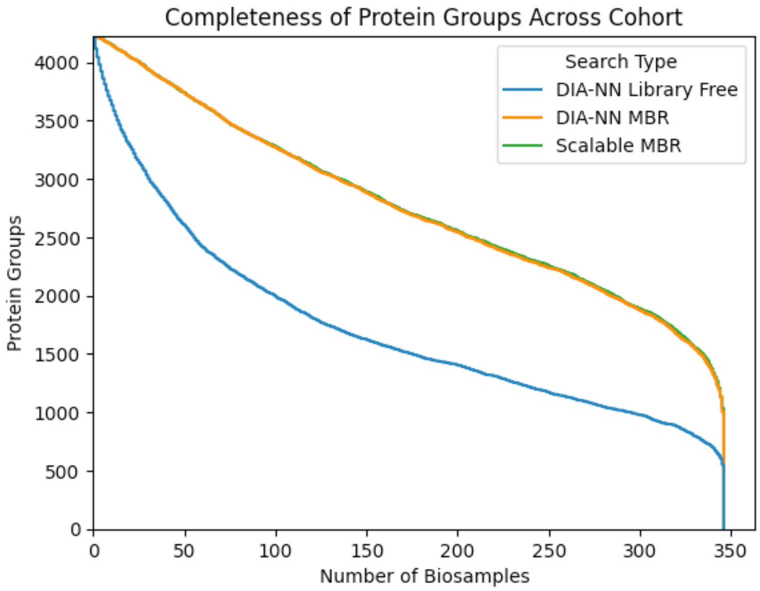


**SFigure 1: MBR increases data completeness across a cohort.** Library free search without match between runs (blue) yields higher missing rate across a cohort. Match between runs increases data completeness to enable better powered studies. Both DIA-NN MBR (orange) and Scalable MBR (green) comparable levels of increased completeness.

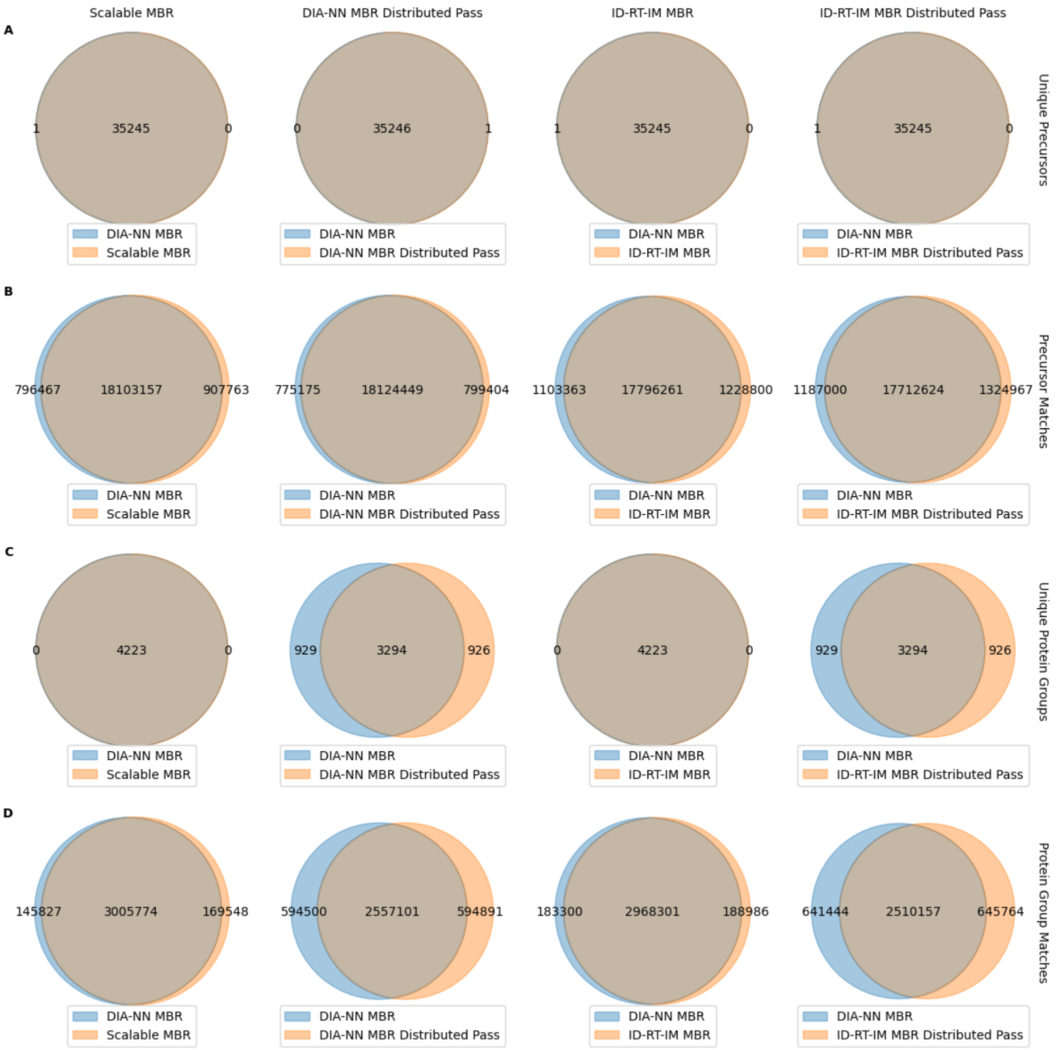


**SFigure 2: Comparison of DIA-NN MBR with alternate MBR Strategies.** Comparison of **A)** unique precursors, **B)** precursor matches, **C)** unique protein groups and **D)** protein group matches. Large differences for the two Distributed Pass alternatives are due to DIA-NN performing a second round of protein inference. Scalable MBR correctly handles modifying the generated library for second pass search without protein re-inference.

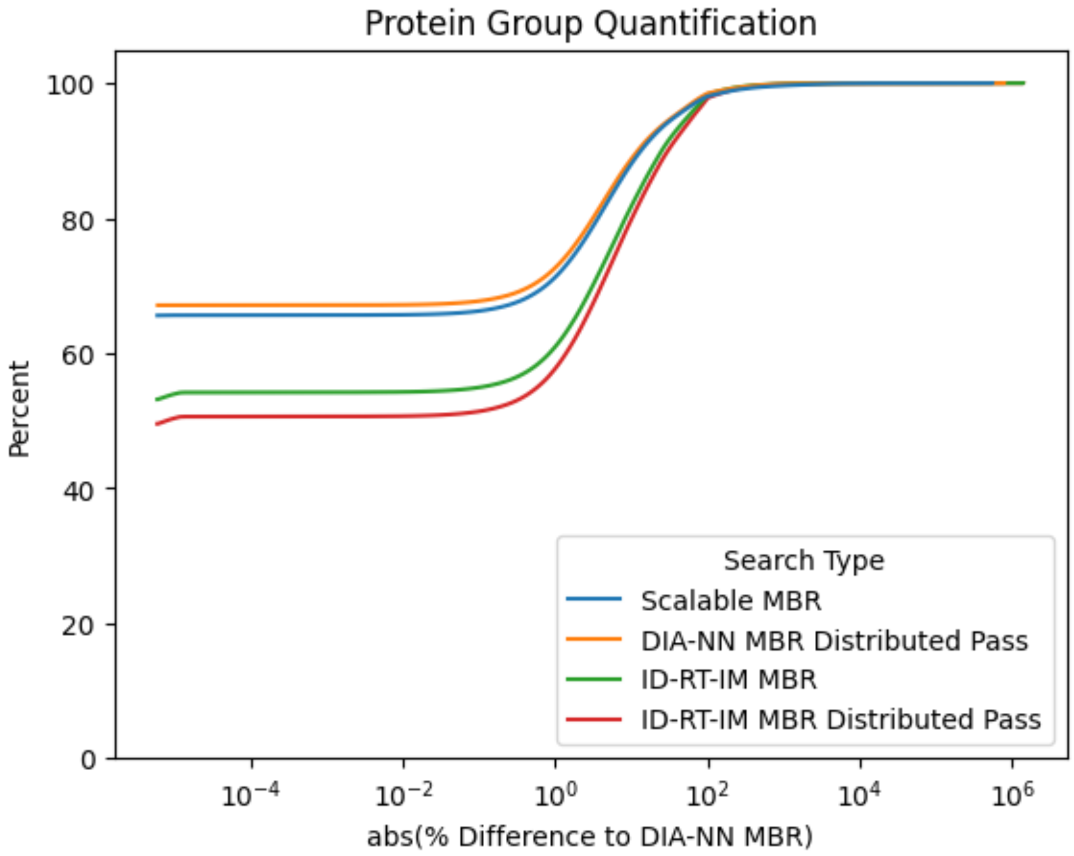


**SFigure 3: Difference in protein group quantification between MBR** strategies. Smart profiling based Scalable MBR more closely matches DIA-NN MBR compared to ID-RT-IM based MBR strategies.
